# BrainQCNet: a Deep Learning attention-based model for the automated detection of artifacts in brain structural MRI scans

**DOI:** 10.1101/2022.03.11.483983

**Authors:** Mélanie Garcia, Nico Dosenbach, Clare Kelly

## Abstract

Analyses of structural MRI (sMRI) data depend on robust upstream data quality control (QC). It is also crucial that researchers seek to retain maximal amounts of data to ensure reproducible, generalizable models and to avoid wasted effort, including that of participants. The time-consuming and difficult task of manual QC evaluation has prompted the development of tools for the automatic assessment of brain sMRI scans. Existing tools have proved particularly valuable in this age of Big Data; as datasets continue to grow, reducing execution time for QC evaluation will be of considerable benefit. The development of Deep Learning (DL) models for artifact detection in structural MRI scans offers a promising avenue toward fast, accurate QC evaluation. In this study, we trained an interpretable Deep Learning model, ProtoPNet, to classify minimally preprocessed 2D slices of scans that had been manually annotated with a refined quality assessment (ABIDE 1; *n* = 980 scans). To evaluate the best model, we applied it to 2141 ABCD scans for which gold-standard manual QC annotations were available. We obtained excellent accuracy: 82.4% for good quality scans (Pass), 91.4% for medium to low quality scans (Fail). Further validation using 799 scans from ABIDE 2 and 750 scans from ADHD-200 confirmed the reliability of our model. Accuracy was comparable to or exceeded that of existing ML models, with fast processing and prediction time (1 min per scan, GPU machine, CUDA-compatible). Our attention model also performs better than traditional DL (i.e., convolutional neural network models) in detecting poor quality scans. To facilitate faster and more accurate QC prediction for the neuroimaging community, we have shared the model that returned the most reliable global quality scores as a BIDS-app (https://github.com/garciaml/BrainQCNet).

## 1. Introduction

Analyses of structural MRI (sMRI) data depend on robust upstream data quality control. This is particularly true for predictive analyses incorporating machine learning techniques, where artifacts and noise may severely bias results and jeopardize generalisability (Backhausen et al., 2016; Gilmore et al., 2019; White et al., 2018; Reuter et al., 2015). Artifacts related to participant motion are a particular concern when working with very young participants, or those with neurodevelopmental diagnoses, such as Autism Spectrum Disorder and Attention-Deficit/Hyperactivity Disorder (Rauch, 2005; Nordahl et al., 2016). In such settings, data collection is usually a demanding and costly task, and it is crucial that researchers retain the maximum amount of usable data to build realistic models.

In this age of big data, manual QC evaluation of sMRI data through visual inspection is a time-consuming and monotonous task, prompting the development of new tools for automatic (full or partial) quality assessment of brain sMRI scans (Esteban et al., 2017; Sujit et al., 2019; Shehzad et al., 2015; Keshavan et al., 2019; White et al., 2018; Alfaro-Almagro et al., 2018; Glasser et al., 2016; Marcus et al., 2013). Such tools typically compute a number of diagnostic metrics using sMRI data to help researchers sort images prior to any analysis (Marcus et al., 2013; Shehzad et al., 2015; Glasser et al., 2016; Esteban et al., 2017; White et al., 2018; Alfaro-Almagro et al., 2018). For example, MRIQC (Esteban et al., 2017) has revolutionized QC of MRI data by providing a reliable and accurate Machine Learning-based assessment of scan quality that <u>has been made freely available</u> to the neuroimaging community as an open-source application. The tool generates 64 image quality metrics, including Contrast to Noise Ratio and Entropy Focus Criterion (Esteban et al., 2017), chosen on the basis of the Preprocessed Connectomes Project (PCP) Quality Assessment Protocol (Shehzad et al., 2015). The MRIQC algorithm uses Machine Learning to find a function that predicts a global quality score for each scan using these metrics. Although highly accessible, automated, and accurate, growth in the size of datasets (e.g., thousands to tens of thousands of sMRI scans for database such as ABCD (Volkow et al., 2018; Karcher and Barch, 2021), ENIGMA (e.g., Whelan et al., 2018) and UK Biobank (Sudlow et al., 2015), prompts a search for developments that can further reduce execution time for QC evaluation. In this study, we evaluate whether Deep Learning models can help advance this goal.

Deep Learning models may prove particularly useful for the task of automated QC. While training a Deep Learning model, such as a convolutional neural network (CNN), may initially take longer than training a traditional Machine Learning (ML) algorithm (because there are more parameters to train), the subsequent processing and inference time is reduced compared to ML (which requires more data preprocessing before inference). This rapid inference makes DL models more scalable for Big Data applications. Studies have already successfully applied DL models to the task of sMRI QC. For example, Sujit et al. (2019) built a CNN model for each axis (sagittal, coronal, axial), and used a fully connected network to return a final prediction based on the intermediary predictions generated by each CNN. Although the model performed well on an multi-site test dataset, it showed poor sensitivity (0.41) when applied to an independent sample. Keshavan et al. (2019) trained a CNN model on slices of scans from a database comprising 200 scans for which expert/gold-standard manual QC was available and 722 scans judged by “citizen scientists.” The AUROC for predicted labels (pass/fail) on a left-out (but non-independent) dataset was 0.99. The authors explained that this high score was due to the fact that the left-out dataset contained scans from similar sites as the training set and the fact that these scans were either very high quality or very low quality, with no intermediate quality scans included in the evaluation. These studies suggest that DL can usefully be applied to predict sMRI scan quality, but highlight the need to ensure that models are generalisable to unseen and independent data that is representative of the range of quality typically observed.

Beyond generalisability, DL models suffer from a lack of interpretability. Visual attention models offer a means to address this. These models mimic human visual attention by identifying the parts of the input image most relevant to the task. For example, when recognising a bird species from a single image, a person might rely on specific details, such as the size, color, or shape of the beak or feathers. Attention-based DL algorithms mimic this process such that the parts of an input that contribute most to prediction (i.e., the most strongly predictive features) can be identified, leading to improved interpretability.

Here, we built on the successes of existing ML and DL approaches and leveraged the advantages of DL attention models to perform automated QC of sMRI data. Specifically, we trained the attention CNN ProtoPNet (Chen et al., 2019), as well as three standard CNNs (VGG19 - (Simonyan and Zisserman, 2015); ResNet152 - (He et al., 2015); DenseNet161 - (Huang et al., 2018)) on 2D slides of sMRI data which had been manually annotated as either good or poor quality. The process used by the ProtoPNet algorithm is similar to the one humans use when we perform manual classification of MRI scans. First, we visually search for the presence of artifacts, slice by slice, in 2D. To judge the quality of a given scan, we focus on specific features in a slice (e.g., the presence of rings or blurring) and compare these features to prototypically corrupted scans. ProtoPNet imitates this human attention process artificially, and returns interpretable output: information about the areas of the input slice identified as being poor quality or defect-free (good). The model also provides another level of interpretability: it points to prototypical cases containing the predictive features.

To train a Deep Learning model, it is crucial that the inputs are correctly labeled. We manually rated 980 structural MRI scans from the ABIDE 1 dataset (Di Martino et al., 2014) guided by (Backhausen et al., 2016), who described four types of artifacts. To train our algorithms, we developed an augmented training set of 270000 2D image slices, derived from 60 scans and a validation set of 1800 2D image slices from 12 scans, perfectly balanced for good quality and very poor quality slices. To identify the best-performing model, we tested the models on the remaining 908 scans from the ABIDE 1 dataset, which had been manually QCed. Finally, we evaluated the best-performing model on independent, multisite datasets: using 2141 scans from ABCD (Volkow et al., 2018; Karcher and Barch, 2021), 799 scans from ABIDE 2 (Di Martino et al., 2017) and 751 scans from ADHD-200 (Bellec et al., 2017).

A key advantage of our algorithm over existing approaches is that it requires only minimal preprocessing, which dramatically reduces the total processing time for every scan (1 minute on a GPU machine, 20 minutes on a CPU machine). Across our independent testing datasets, we observed excellent accuracy that matched or surpassed existing automated QC algorithms. In the context of the growth of Open Science datasets to tens of thousands of participants, our method could offer substantial savings in terms of time and computational resources.

To facilitate fast and accurate QC prediction for the neuroimaging community, we have shared the model that returned the most reliable global quality scores, local predictions of quality, and maps and prototypes of local artifacts as a BIDS-app (https://github.com/garciaml/BrainQCNet). For the fastest performance, we recommend using the GPU version of our app.

## 2. Materials and Methods

### 2.1 Datasets

In our study, we used structural MRI data from <u>ABIDE 1</u> (Di Martino et al., 2014), <u>ABIDE 2</u> (Di Martino et al., 2017), <u>ADHD-200</u> (Bellec et al., 2017) and <u>ABCD</u> (Volkow et al., 2018; Karcher and Barch, 2021). Details of each of the datasets used are provided in **Figure 1**.

**Figure 1.**
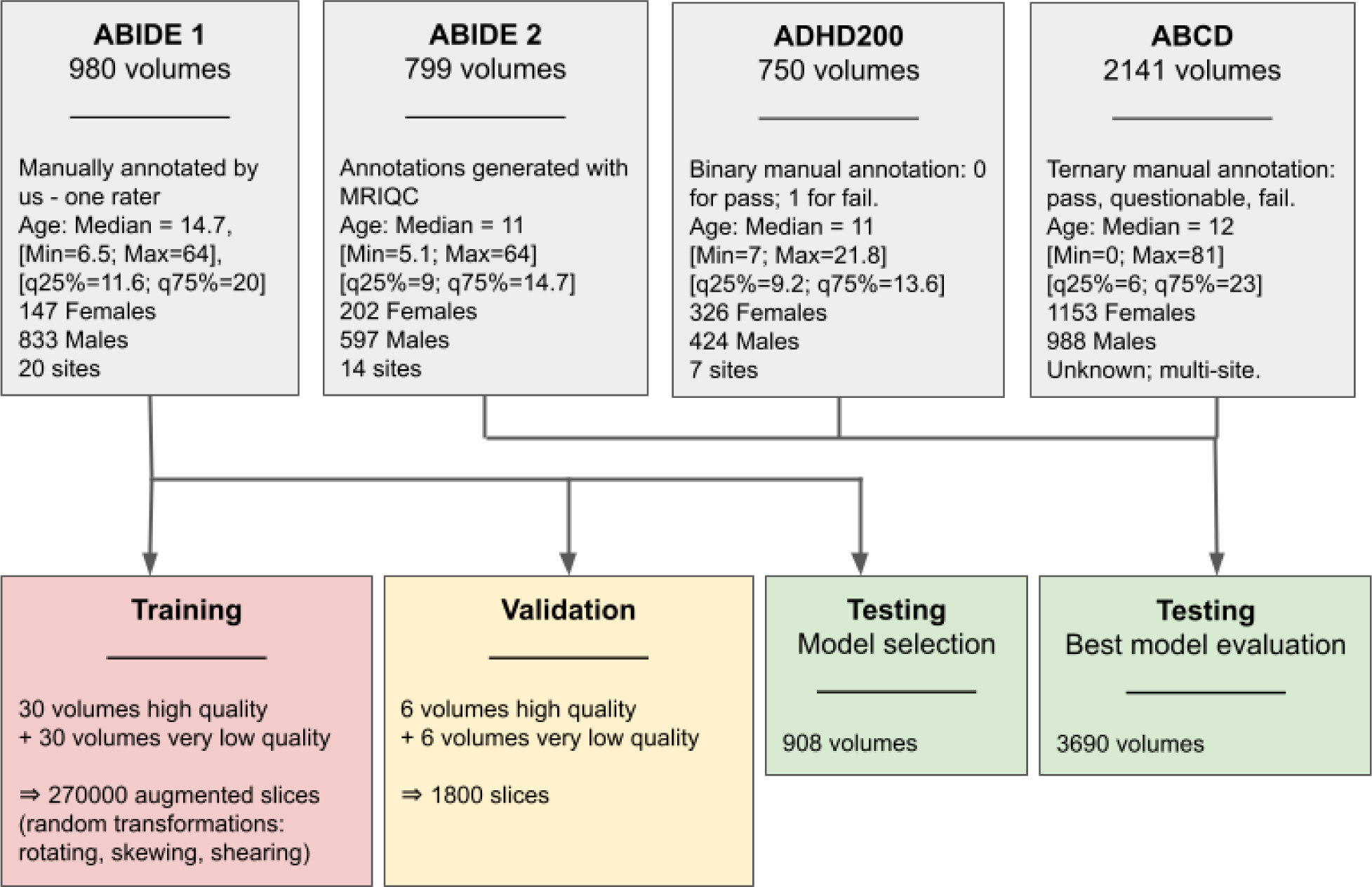
Dataset descriptions and division into training, validation, and testing sets.

### 2.2 Ethics statement

The three databases used in the project - ABIDE 1, ABIDE 2, ADHD200 - are shared by the International Neuroimaging Data-sharing Initiative (http://fcon_1000.projects.nitrc.org/). Each dataset was fully de-identified and anonymized in accordance with the US Health Insurance Portability and Accountability Act (HIPAA). All the datasets were collected and shared in accordance with the local regulations on ethics and data protection. Data usage is unrestricted for non-commercial research purposes; it is openly shared with the scientific community under the license Creative Commons BY-NC-SA. Our work with these open data is approved by the Research Ethics Committee of the School of Psychology at Trinity College Dublin.

Data from the ABCD study were fully de-identified and anonymized, and each data-collecting site obtained informed consent from participants and their parents/guardians. The ABCD study developed guidelines for ethical considerations to be applied by each data-collecting site, and organized a hierarchy of workgroups who assessed whether each step of the collection process conformed to the ABCD guidelines (Clark et al., 2018). Data from the ABCD study were used under a Data Agreement between Trinity College Dublin and Washington University.

### 2.3 Manual Quality Control

One rater (MG) manually annotated 980 MRI scans from ABIDE 1. The annotation was guided by the work of Backhausen et al., (2016), which specified four different types of artifacts: (1) blurring (global or local), (2) ringing, (3) low contrast noise ratio between gray matter and white matter, and (4) low contrast noise ratio (CNR) of subcortical structures. For further details of the artifacts, please see the Supplementary Materials of Backhausen et al., (2016). For each scan and each artifact type, a score between 1 and 4 was given, such that a score of 1 indicates absence of that artifact while scores of 2, 3, and 4 indicate the presence of that artifact at worsening degrees of severity (where 4 is the worst).

For each 3D scan, we also noted whether each of the four artifacts was evident either locally or globally. When no artifact was observed (score = 1,1,1,1), we labeled the 3D scan as good quality (Class 0). Otherwise, we labeled the 3D scan as poor quality (Class 1; see **Figure 2**). Class 1 is a wide spectrum that includes scans with localized artifacts (e.g., score = 1,2,2,1) as well as very low quality, globally disrupted scans (score = 4,4,4,4 and artifacts present on all the slices of the volume). These labels - Class 0 and Class 1 - were used as the true values on which our models were trained and tested.

**Figure 2.**
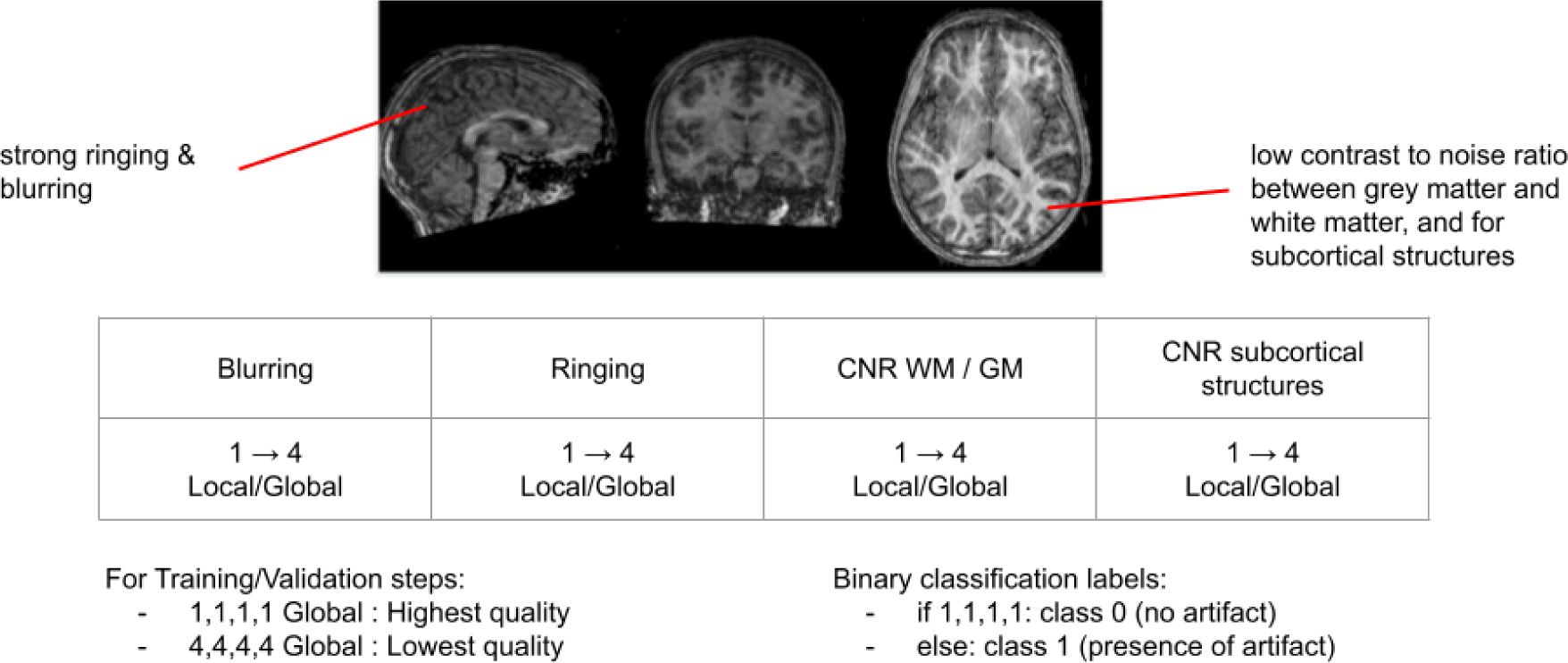
Description of our system for manual sMRI scan quality annotation.

### 2.4 Training & Validation Datasets

To create a set of images on which to train our Deep Learning algorithm, we identified 30 high quality scans (randomly selected from those labeled Class 0) and 30 highly corrupted/poor quality scans (randomly selected from all the scans labeled Class 1 and scored 4,4,4,4) from the 980 ABIDE 1 scans we had manually annotated. We also created a within-training validation set comprising 6 further high quality Class 0 scans and 6 very low quality Class 1 (i.e. score=4,4,4,4 and artifact present on all the slices) scans. Importantly, these training and validation sets included all the highly corrupted scans (i.e., score=4,4,4,4). We did this to provide a balanced training (same number of Class 1 and Class 0 scans) and to maximize the chances of obtaining meaningful prototypes representative of scan artifacts and corruption.

Chen et al. (2019) found that the ProtoPNet algorithm worked better on cropped images, so each 3D scan was tightly cropped to remove empty space, then converted from Nifti format to 2D PNG images (using Med2Image https://github.com/FNNDSC/med2image). For each scan there were between 150-200 2D slices for each of the 3 orientations (sagittal, coronal, axial); resulting in approximately 450-600 images per scan. The first and last 20 slices of each image stack were discarded since they contained little brain tissue. Taking a random sample of 50 slices per axis, per scan, we created a training set comprising 4500 high quality and 4500 poor quality 2D slices from all the 60 scans in the training set. A validation set of 1800 slices, also balanced for quality, was created in the same way.

Next, the training set was augmented with a set of random transformations (using the library Augmentor https://github.com/mdbloice/Augmentor) which rotated, skewed, and sheared the images. This yielded an augmented training set of 270000 images. Data augmentation is used to prevent overfitting in Deep Learning, thus improving generalizability of the algorithms.

All 2D images from good quality scans (Class 0) were defined as Label 0 and all 2D images from poor quality scans (Class 1) were defined as Label 1. The algorithm was trained to perform a binary classification between Label 0 and Label 1 2D slices using the augmented training set (*n* = 270000 slices), and validation accuracy was computed every 2 epochs (n = 1800 slices). An epoch is a hyperparameter that defines the number of times that the learning algorithm has optimized the parameters on the entire training dataset. This process of data preparation, training, and validation is summarized in **Figure 1**.

Since predictions were performed at the level of slices, to generate a global prediction for each scan, we computed the proportion of slices with a prediction of Label 1 (poor quality) and applied a threshold of 0.5. If greater than 50% of slices for a given scan were predicted Label 1, the entire scan was classified as Class 1 (poor quality). Below this threshold, the entire scan was classified Class 0 (good quality). We note that this is an arbitrary threshold and that different thresholds may be preferable, depending on the particular goal of subsequent analyses. Our BIDS-app (https://github.com/garciaml/BrainQCNet) returns a CSV file containing scan identifiers and probability scores, allowing for the specification of a new threshold for tailored scan classification.

### 2.5 Testing Set for Model Selection

To identify the best-performing model (see **Section 3.3**), we generated predictions for the remaining 908 scans from ABIDE 1 (Di Martino et al., 2014), which we had manually annotated. For each scan, 450-600 2D slice images were created using the process described above (**Section 2.4**).

### 2.6 Independent Testing Sets for Evaluation

After identifying the best-performing model, we performed an evaluation using independent testing sets comprising 2D slice images created using the process described above, for 3690 sMRI scans obtained from the following sources (see **Figure 1**):

- 2141 scans from ABCD (Volkow et al., 2018; Karcher and Barch, 2021). These scans had been manually QC’ed by two or more reviewers (Hagler et al., 2019), following the recommendation from the ABCD Data Analytics and Informatics Core (DAIC) (Saragosa-Harris et al., 2022), with ternary classification: pass, questionable, fail;
- 799 scans from ABIDE 2 (Di Martino et al., 2017) with QC classification generated by the MRIQC algorithm (see **Section 2.8**, below);
- 750 scans from ADHD-200 (Bellec et al., 2017). These scans had been manually QC’ed by 1 or 2 human raters (Bellec et al., 2017) with binary classification: pass, fail.

### 2.7 Deep Learning Algorithm

The algorithm we used, ProtoPNet (Chen et al., 2019), is a Deep Learning Attention model that reproduces the human manual process for classifying images. The network consists of a regular convolutional neural network, followed by a prototype layer and a fully connected layer with weight matrix and no bias. Here, we compared three different architectures for the regular convolutional network: VGG19 (Simonyan and Zisserman, 2015), ResNet152 (He et al., 2015) and DenseNet161 (Huang et al., 2018). These three models are well known Deep Learning algorithms for image classification, and have shown good performance for 2D images (Simonyan and Zisserman, 2015; He et al., 2015; Huang et al., 2018). In Machine Learning, it is common to compare different types of algorithm for a given problem, to detect overfitting and to identify the best-performing algorithm (Hastie et al., 2009).

In their approach, (Chen et al., 2019) constrained each convolutional filter to be identical to a latent training patch, to make every convolutional filter interpretable as visualisable prototypical image parts. In our study, the “prototypes” or “prototypical images” corresponded to the Class 0 (good quality) and Class 1 (poor quality) images of the augmented training set. The algorithm works, in part, by comparing images in the validation and test sets to parts of the prototypes. The number of images selected randomly as prototypes during each epoch of training was set to 2000.

In the ProtoPNet global architecture, the prototype layer computes similarity scores between the convolutional filters of the input image and the ones from the 2000 prototypes at a fixed epoch.The similarity scores are computed with an inverted L2 norm distance.

(Chen et al., 2019) explained that given a convolutional output *z* = *f*(*x*), the j-th prototype unit *g_p_j__* in the prototype layer *g_p_* computes the squared *L^2^* distances between the j-th prototype *p*_*j*_ and all patches of *z* that have the same shape as *p*_*j*_, and inverts the distances into similarity scores. The result is an activation map of similarity scores whose value indicates the strength of similarity between the input image and a prototype.

Mathematically, the prototype unit *g_p_j__* computes 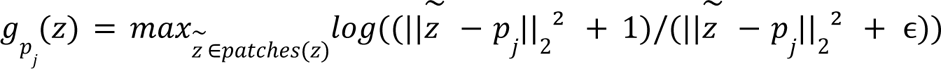. The function *g_p_j__* is monotonically decreasing with respect to 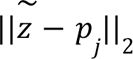 (if 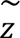 is the closest latent patch to *p_j_*). If the output of the j-th prototype unit *g_p_j__* is large, then there is a patch in the convolutional output that is (in 2-norm) very close to the j-th prototype in the latent space, and this in turn means that there is a patch in the input image that has a similar concept to what the j-th prototype represents.

Next, the fully connected layer predicts the label of the input image from the 2000 similarity scores. We obtained probability scores by applying the softmax function to the output logits of the fully connected layer. In theory, this method of regularization and comparison should improve the generalizability of the algorithm. More mathematical details of the ProtoPNet model are given in (Chen et al., 2019); **Figure 3(b)** illustrates its architecture in our context.

**Figure 3.**
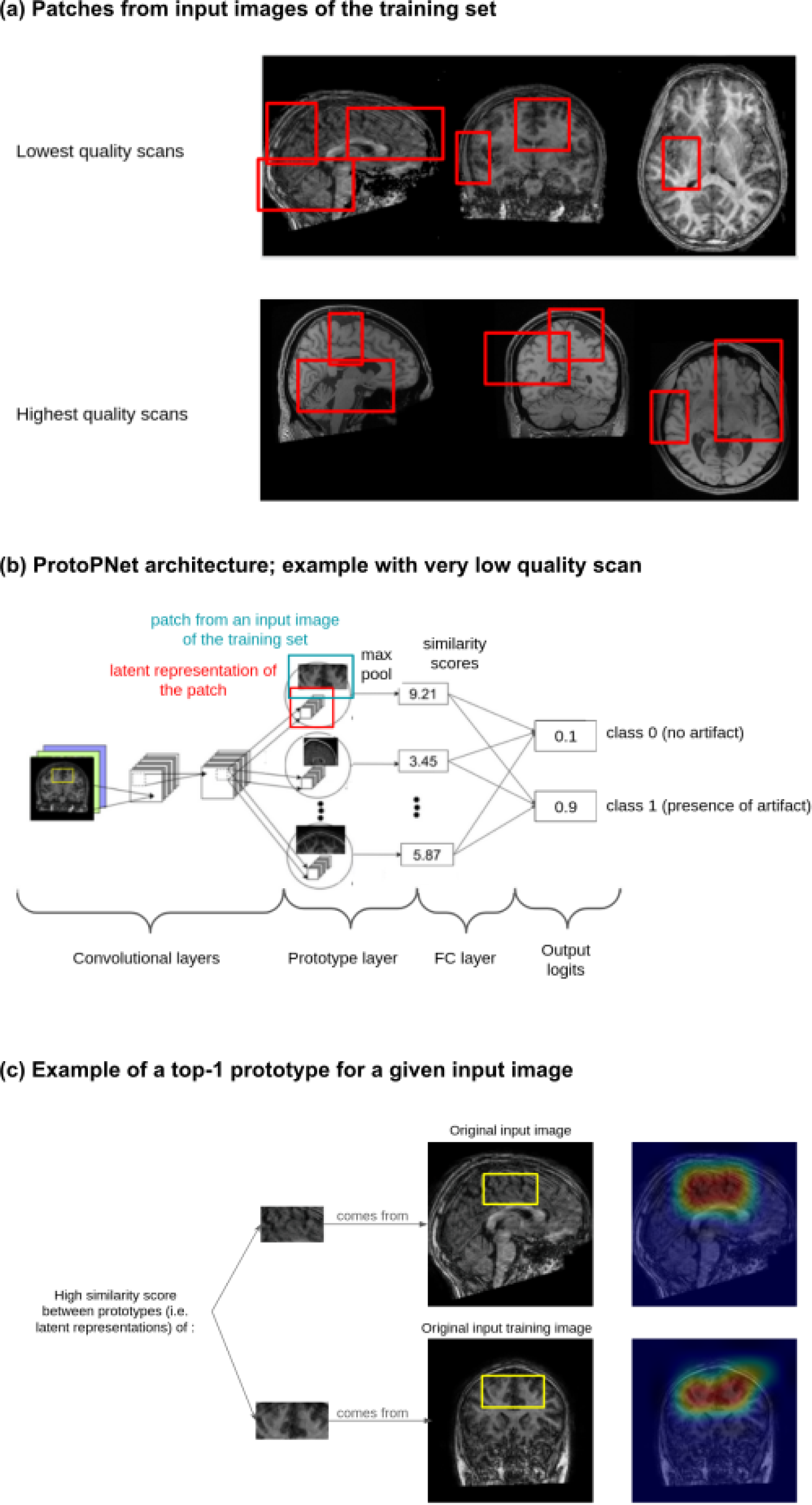
The ProtoPNet approach for automatic QC of brain sMRI scans. (a) Patches taken from input 2D slices of the training set; (b) Architecture of the ProtoPNet model; (c) Example of a top-1 prototype (i.e., the prototype from the training set with the highest score for similarity with the input patch) for a given input 2D slice.

We initiated training using ImageNet (Deng et al., 2009), drawn from the model zoo of Pytorch (https://pytorch.org/serve/model_zoo.html). We used the same initialisation parameters as previous experiments (Chen et al., 2019), including 5 “warming” epochs for which no accuracy was computed (where each epoch is a step during which the algorithm is optimized by all the images of the training set). Because of the GPU memory demands of this process, optimization is achieved iteratively using small batches of data. Here, we used the same batch sizes as (Chen et al.; 2019): 80 for the training and 100 for the testing phase. During training time, we validated every 2 epochs by assessing the prediction accuracy of the model for slices from the scans in the validation set.

We trained our models in a distributed way on AWS cloud instances of type p3.8xlarge and p3.16xlarge initialized with the AMI Deep Learning. The instances correspond to 4 or 8 GPUs NVIDIA V100. We trained ResNet152 on 20 epochs and VGG19 and DenseNet161 on 30 epochs. We saved models and associated prototypes every 10 epochs.

### 2.8 MRIQC

MRIQC (Esteban et al., 2017) was conceived as a tool to permit more reliable and efficient QA/QC of MRI data through visual reports. It integrates a classifier to provide an automatic assessment of the quality of brain structural and functional MRI scans. The MRIQC classifier is based on a Machine Learning algorithm that was trained on a large number of metrics of quality previously extracted and computed from raw scans. As outlined in the introduction, these metrics were chosen as part of the Preprocessed Connectomes Project (PCP) Quality Assessment Protocol (Shehzad et al., 2015) to harmonize the assessment of the quality of brain MRI scans (Shehzad et al., 2015), like the signal-to-noise ratio. The output of MRIQC is a score and a binary prediction (pass/fail) for each scan.

This method is reliable (accuracy estimated to 76%±13% on new sites, using leave-one-site-out cross-validation, accuracy of 76% on a held-out dataset of 265 scans; Esteban et al., 2017), and widely employed.

Here, we used the MRIQC classifier to generate predictions of the quality of each scan on ABIDE 2 (Di Martino et al., 2017; 799 scans). We used the default MRIQC threshold for classification. In particular, we used the BIDS-app poldracklab/mriqc:0.9.6 (on DockerHub) to run the MRIQC classifier as is. We treated these MRIQC-based predictions as the “ground truth” against which we compared the results of our algorithm.

We also compared the distribution of the scores returned by MRIQC for ABIDE 1 (n = 980 scans; Di Martino et al., 2014) with the distribution of scores returned by our models. In particular, we examined the discrimination between good quality scans (score=1,1,1,1) and medium quality (artifacts present only locally on the volume and/or medium intensity artifacts) and low quality ones (score=4,4,4,4 and artifacts present on all the slices of all the volume).

### 2.9 Comparison with Traditional CNN Models

To provide a comprehensive evaluation of the attention model (ProtoPNet) approach, we also built three traditional CNN models for comparison. To do this, we used the pre-trained CNN models, VGG19, ResNet152, DenseNet161, drawn from the model zoo of Pytorch (https://pytorch.org/serve/model_zoo.html). We used the same training and validation sets, learning parameters, and methods described above.

### 2.10 Data and Code Availability

Three of the datasets used in the project - ABIDE 1, ABIDE 2, ADHD200 - are openly shared by the International Neuroimaging Data-sharing Initiative (http://fcon_1000.projects.nitrc.org/). Access to ABCD data is available upon request (https://nda.nih.gov/abcd/request-access).

All global predictions of quality for the 4670 scans we used from the ABIDE 1 & 2, ADHD200 and ABCD databases are available through the GitHub repository: https://github.com/garciaml/BrainQCNet_paper_results.

To maximize the reproducibility of our analyses and usability of our model, all the code to build the BIDS-apps is available on two other GitHub repositories (https://github.com/garciaml/BrainQCNet_CPU for users of CPU machines and https://github.com/garciaml/BrainQCNet_GPU for users of GPU machines compatible with CUDA technology). Non-containerized version for CPU is also available (https://github.com/garciaml/BrainQCNet_CPU_non_containerized).

We have integrated the best-performing QC model into an open-source BIDS-app (Gorgolewski et al., 2017), to share it with the neuroimaging community in a ready-to-use format. Documentation for our BIDS-app for CPU or GPU is available here: https://github.com/garciaml/BrainQCNet. We have also shared our trained CNN baseline models for reuse: https://github.com/garciaml/BrainQCNet_CNN_GPU.

The following BIDS-apps are available on DockerHub:

- garciaml/brainqcnet-cnn: the best CNN model (which provides a control/comparison for the model based on ProtoPNet architecture);
- garciaml/bids-pytorch-cuda: a template for Deep Learning BIDS-app running on GPU/CUDA machines using the Pytorch framework;
- garciaml/brainqcnet: the best-performing model identified in this study, for use on GPU/CUDA machines;
- garciaml/brainqcnetcpu: the best-performing model of this study, for us on CPU machines.

Our apps and code are available under the Apache License, Version 2.0, January 2004.

We have also created and shared two demo videos explaining how to run our app on CPU and on GPU machines compatible with CUDA technology (links available on https://github.com/garciaml/BrainQCNet).

## 3. Results

### 3.1 Annotations

Manual QC inspection of 980 scans from ABIDE 1 (Di Martino et al., 2014) identified 564 high quality scans (Class 0), 36 very low quality scans (i.e. globally corrupted and score=4,4,4,4; which we used in the training and validation sets), and 380 scans with either local artifacts or with mild-moderate global corruption. Local ringing (likely reflecting motion) was the most commonly occurring local artifact, and was often combined with other artifact types.

### 3.2 Training performance

In the results and figures below, we use the following naming convention: the prefix “proto-” corresponds to the ProtoPNet algorithm, while the suffix indicates the CNN architecture: V19 for VGG19, R152 for ResNet152, or D161 for DenseNet161 (see **Section 2.7**).

We obtained excellent accuracy for the detection of good (Class 0) and bad (Class 1) quality slices during training. From epoch 10, accuracy for the three attention models - proto-V19, proto-R152, proto-D161, was above 99% on the Training set and above 95% on the Validation set. This means that more than 99% of the 270000 training images were accurately classified from epoch 10. Likewise, more than 95% of the 1800 validation slices were accurately classified from epoch 10. Looking at performance on the validation set, the model proto-D161 out-performed proto-V19 and proto-R152 (see **Figure 4, left**).

**Figure 4.**
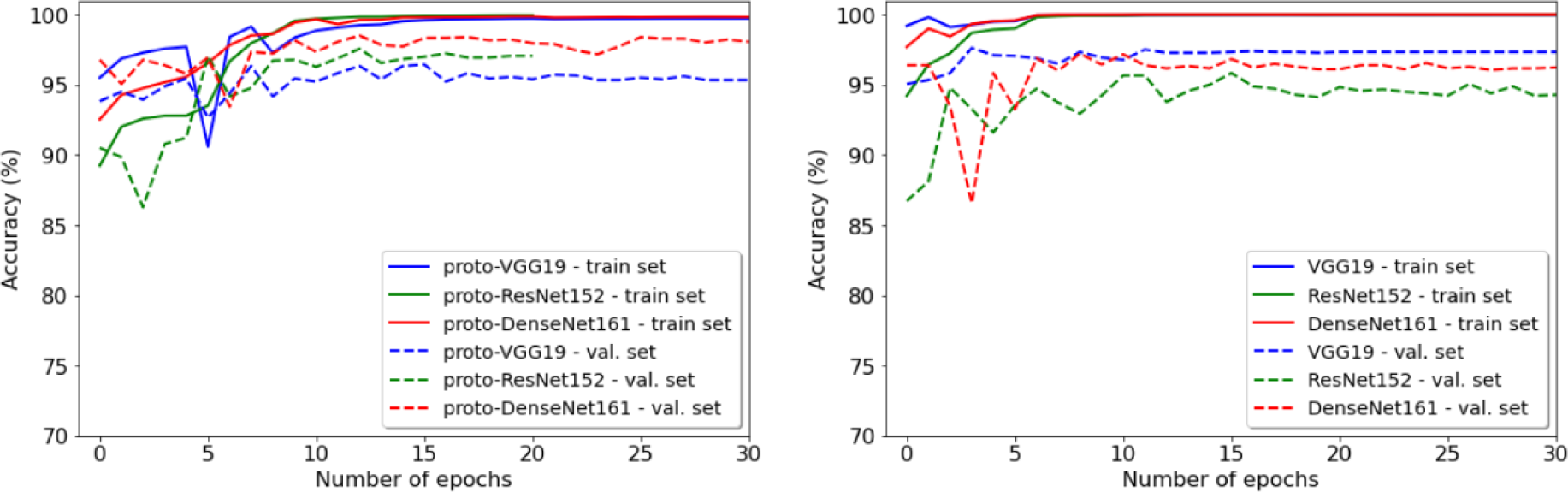
Evolution of accuracy across epochs for the Training and Validation sets; (left) training performance of the ProtoPNet models; (right) training performance of the traditional CNN models.

The traditional CNN comparator models also converged quickly (see **Figure 4, right**). The CNN models (VGG19, ResNet152, DenseNet161) trained on 15 epochs were used as comparators for the main attention models (proto-V19, proto-R152, proto-D161) in all further analyses.

### 3.3 Selecting the best model using ABIDE 1

As described above (Section 2.4), predictions (Class 0/1) were performed at the level of 2D slices from a given scan. To generate a global prediction for each scan, we applied a threshold such that if >50% of slices for a given scan were predicted Label 1, the entire scan was classified as Class 1 (poor quality). Below this threshold, the entire scan was classified Class 0 (good quality). Producing a binary scan-level class prediction is useful in the QC context, because it provides a pass (Class 0) or fail (Class 1) outcome. However, there are likely to be applications for which an examination of the value of the proportion itself might be warranted, since this value gives more information about the quality of the scan. In analyses and comparisons performed below, we have operationalised this proportion as a probability - specifically, it is the frequentist probability that a given scan is corrupted by an artifact. Similarly, there will be applications where a different threshold (e.g., >0.4 = Class 1) may be preferable, depending on the particular goal of subsequent analyses. Our BIDS-app (https://github.com/garciaml/BrainQCNet) allows for the specification of a threshold for scan classification.

**Table 2** compares the specificity and sensitivity scores for each model. While specificity is very high (>95%) for all the models (with the exception of MRIQC = 91.1%), sensitivity is relatively low. The highest sensitivity is achieved by the model proto-R152 trained on 10 epochs (47.89%) followed by the MRIQC classifier (41.58%). This may be explained by the fact that since the most severely corrupted scans were used for training, the Test set contains scans that are generally of lower and more variable severity of artifact and poor quality. Scans of moderate quality (less severe global artifact, or very localized artifact) likely yield probabilities between 0.4 and 0.5. This means that the Class predicted is 0 (good quality), the scan is of moderate rather than high quality. **Supplemental Figure S2** shows the distribution of probabilities for each model and each dataset.

**Table 2.** Accuracy (Acc.) and ROC AUC (AUC) scores for Training, Validation, and Test sets. Specificity (“Sp.”) and Sensitivity (“Sens.”) scores on the testing set. For each of the attention models, performance after 10, 20, and 30 training epochs (parameter optimization steps) is shown.

| Model |  | Training<br>(60 scans) | Validation<br>(12 scans) | Test<br>(908 scans) |  |  |
| --- | --- | --- | --- | --- | --- | --- |
|  |  |  |  | All Scans | Artifact-free<br>Class 0<br>(528 scans) | With Artifact<br>Class 1<br>(380 scans) |
| **proto-D161**<br>**epochs** | **10** | Acc. = 100%<br>AUC = 1 | Acc. = 100%<br>AUC = 1 | Acc. = 69.8%<br>AUC = 0.775 | Sp. = 99.4% | Sens. = 28.7% |
| **proto-D161**<br>**epochs** | **20** | Acc. = 100%<br>AUC = 1 | Acc. = 100%<br>AUC = 1 | Acc. = 64.7%<br>AUC = 0.774 | **Sp. = 100%** | Sens. = 15.5% |
| **proto-D161**<br>**epochs** | **30** | Acc. = 100%<br>AUC = 1 | Acc. = 100%<br>AUC = 1 | Acc. = 62%<br>AUC = 0.758 | **Sp. = 100%** | Sens. = 9.2% |
| **proto-R152**<br>**epochs** | **10** | Acc. = 100%<br>AUC = 1 | Acc. = 100%<br>AUC = 1 | **Acc. = 75.4%**<br>**AUC = 0.825** | Sp. = 95.3% | **Sens. = 47.9%** |
| **proto-R152**<br>**epochs** | **20** | Acc. = 100%<br>AUC = 1 | Acc. = 100%<br>AUC = 1 | Acc. = 68.7%<br>AUC = 0.811 | Sp. = 99.6% | Sens. = 25.8% |
| **proto-V19**<br>**10 epochs** |  | Acc. = 100%<br>AUC = 1 | Acc. = 100%<br>AUC = 1 | Acc. = 67.2%<br>AUC = 0.823 | Sp. = 99.6% | Sens. = 22.1% |
| **proto-V19**<br>**20 epochs** |  | Acc. = 100%<br>AUC = 1 | Acc. = 100%<br>AUC = 1 | Acc. = 70.0%<br>AUC = 0.849 | Sp. = 99.1% | Sens. = 29.7% |
| **proto-V19**<br>**30 epochs** |  | Acc. = 100%<br>AUC = 1 | Acc. = 100%<br>AUC = 1 | Acc. = 71.8%<br>AUC = 0.847 | Sp. = 98.5% | Sens. = 34.7% |
| **MRIQC_CLF** |  | Acc. = 96.7%<br>AUC = 0.767 | Acc. = 100%<br>AUC = 1 | Acc. = 70.4%<br>AUC = 0.724 | Sp. = 91.1% | Sens. = 41.6% |
| **CNN-DenseNet**<br>**161**<br>**15epochs** |  | Acc. = 100%<br>AUC = 1 | Acc. = 100%<br>AUC = 1 | Acc. = 68.1%<br>AUC = 0.787 | Sp. = 99.6% | Sens. = 24.2% |
| **CNN-ResNet15**<br>**2**<br>**15 epochs** |  | Acc. = 100%<br>AUC = 1 | Acc. = 100%<br>AUC = 1 | Acc. = 69.3%<br>AUC = 0.792 | Sp. = 99.4% | Sens. = 27.4% |
| **CNN-VGG19**<br>**15 epochs** |  | Acc. = 100%<br>AUC = 1 | Acc. = 100%<br>AUC = 1 | Acc. = 68.6%<br>AUC = 0.781 | Sp. = 99.6% | Sens. = 25.5% |

**Table 2** compares the classification accuracies for global quality of the Training, Validation, and Test sets, obtained for each of the models, including MRIQC and the CNN models. These results show that the best model for the prediction of sMRI scan global quality is proto-R152 trained on 10 epochs. This model is at least as accurate as MRIQC and the CNN models. **Supplemental Figures S1 and S2** provide further illustrations of the distribution of probability scores across models.

We identified proto-R152 (after 10 epochs) as the best model among those compared. **Supplemental Figure S3** shows the distributions of probability scores for the proto-R152 model for ABIDE 1 scans with different types/levels of severity of artifact.

As described above, each algorithm selected 2000 prototype images from the augmented training set of 270000 images during each training epoch. **Figure 3** and **Supplemental Figures S5 and S6** provide examples of the prototypes. Examination of the prototypes for proto-R152 after 10 epochs suggested a set of diverse prototypes that were highly relevant for the type of artifacts detected in the ABIDE I dataset.

Further, the distribution of accuracies across categories and sites does not appear to suggest a site effect (see **Supplemental Table S1**), and there was no difference in the global distribution of probabilities between the three axes (sagittal, coronal, axial).

### 3.4 Evaluation using ABCD (2141 scans)

The ABCD dataset was annotated with gold-standard manual QC judgments thanks to the workgroups performing data collection and quality control (Karcher and Barch, 2021). We tested our algorithm on 2141 of these manually QCed scans. **Figure 5** compares the distribution of probabilities between QC categories (pass, questionable, fail) for these 2141 ABCD scans, computed by the best-performing model (proto-R152 trained on 10 epochs). It shows that, although there is some overlap, the central tendency and distribution of probability scores differ between pass and fail categories. There is greater overlap between scores of the questionable and pass categories, which is to be expected. We confirmed this observation by performing Mann-Whitney U-tests (because the normality assumption for a T-test was not verified for any of the samples; see **Supplemental Table S2)**.

**Figure 5.**
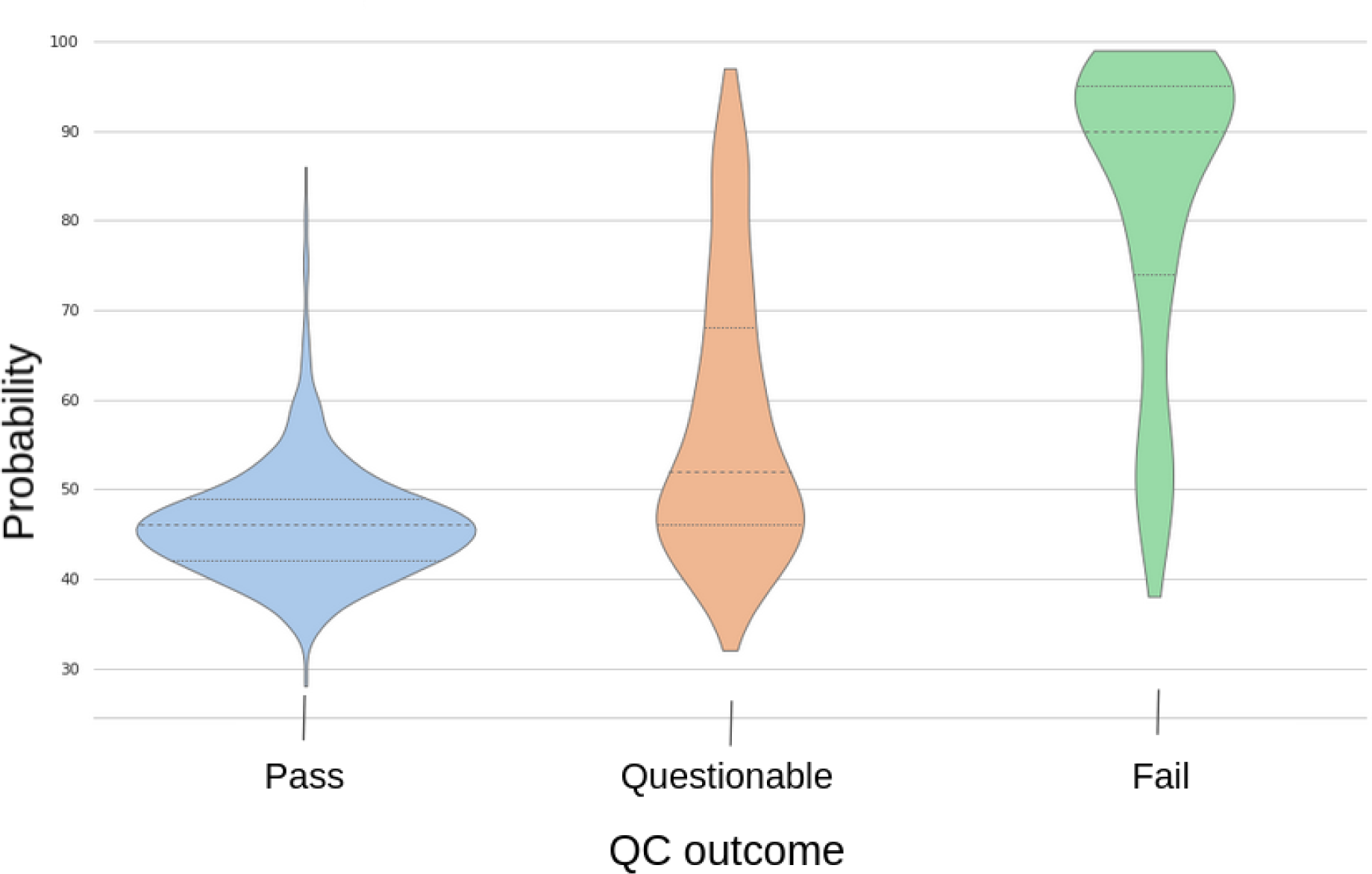
The distribution of probabilities between the true QC categories (pass, questionable, fail) for ABCD data (2141 scans), computed by proto-R152 trained on 10 epochs.

**Table 3** shows that our algorithm showed better accuracy for the category “fail” than the comparison models. Conversely, the three CNN baseline models and MRIQC (tested on 410 of the 2141 scans, due to the time required for processing) initially performed better than proto-R152 when predicting the category “pass”. Upon closer inspection, we found that 311 “pass” scans had probabilities between 0.5 and 0.6. When these scans are removed and only scans with probabilities lower than 0.5 or greater than 0.6 are retained, accuracy was 96.4% for the pass category. It is possible that our algorithm detected mild artifacts that were not considered significant by human raters. Accordingly, depending on the application, we suggest a second verification - either manual checking or a second model - for scans with “borderline” probabilities (0.5-0.6).

**Table 3.**
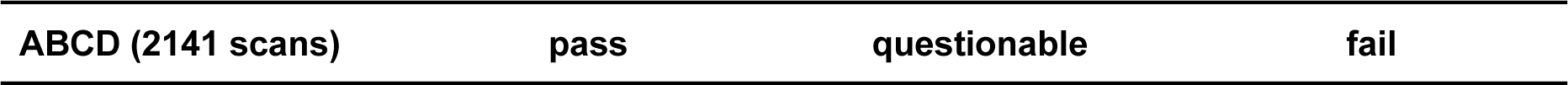
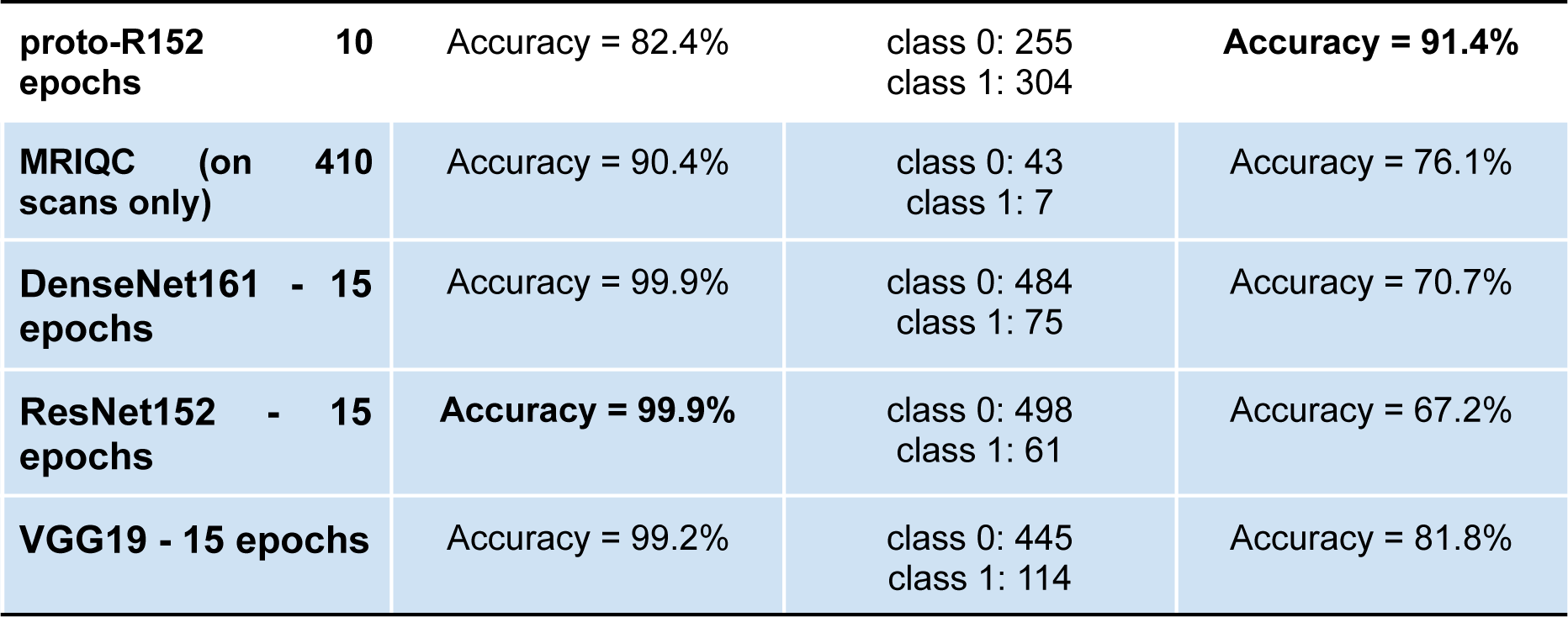
Accuracy of predictions for each of the manually determined QC categories (pass, questionable, fail) for ABCD data (2141 scans).

### 3.5. Evaluation using ABIDE 2 (799 scans) and ADHD-200 (750 scans)

To further evaluate our tool using independent data, we ran the MRIQC classifier on 799 scans from the ABIDE 2 dataset and treated its predictions as ground truth. The MRIQC classifier predicted 588 Class 0 (pass) scans and 211 Class 1 (fail). Accuracy for our proto-R152 was 75.5%. The ROC AUC score was 0.72.

We also evaluate our model using the ADHD200 dataset, which includes manual QC (pass, fail) annotations for 750 scans. Our proto-R152 model attained an accuracy score of 79.2% and a ROC AUC score of 0.76. Sensitivity was greater than for the CNN baseline models but specificity was lower. These results are summarized in **Table 4**.

**Table 4.** Accuracy (“Acc.”), ROC AUC (“AUC”), Specificity (“Sp.”) and Sensitivity (“Sens.”) scores for the proto-R152 and CNN comparison models for ABIDE 2 (true quality annotations obtained by the predictions of the MRIQC classifier) and ADHD200.

| ABIDE 2 - QC prediction by MRIQC |  |  |  | ADHD200 |  |  |
| --- | --- | --- | --- | --- | --- | --- |
|  | All | 588<br>uncorrupt<br>ed scans -<br>class 0 | 211<br>corrupted<br>scans -<br>class 1 | All | 711<br>uncorrupted<br>scans - class<br>0 | 39 corrupted<br>scans - class<br>1 |
| **proto-R152**<br>**10 epochs** | Acc. = 75.5%<br>AUC = 0.718 | Sp. =<br>83.5% | **Sens. =**<br>**53.1%** | Acc. = 79.2%<br>**AUC = 0.76** | Sp. = 80.2% | **Sens. = 61.5%** |
| **DenseNet161**<br>**15 epochs** | **Acc. = 80.1%**<br>AUC = 0.726 | Sp. =<br>94.6% | Sens. =<br>39.8% | **Acc. = 90.0%**<br>AUC = 0.747 | **Sp. = 92.4%** | Sens. = 46.2% |
| **ResNet152**<br>**15 epochs** | Acc. = 79.8%<br>**AUC = 0.742** | Sp. =<br>93.7% | Sens. =<br>41.2% | Acc. = 88.4%<br>AUC = 0.674 | Sp. = 90.9% | Sens. = 43.6% |
| **VGG19**<br>**15 epochs** | Acc. = 79.5%<br>AUC = 0.679 | Sp. =<br>**94.7%** | Sens. =<br>37.0% | Acc. = 89.3%<br>AUC = 0.696 | Sp. = 91.6% | Sens. = 48.7% |

### 3.6. Model Interpretability

What features of the input data does our model rely on for prediction? This question relates to the interpretability of the model, which is often challenging for Deep Learning models, relatively to conventional Machine Learning methods. Interpretability is important, not only for revealing the input features that contribute most to classification, but also for pointing to opportunities for model improvement.

First, we considered the prototypes (the 2000 images from the augmented training set of 270000 images selected during each training epoch) used by the attention models (proto-V19, proto-R152, proto-D161) and assessed whether these were well balanced in terms of the types of artifacts represented. We identified the top 5 prototypes (i.e. the 5 prototypes with the highest similarity scores with patches of 2D input slices) for each of the three axes (axial, sagittal, coronal) and observed that two prototypes (ringing and blurring) were highly prevalent among the top 5 (**Figure S4).** We observed that the prototypes used by the best-performing model, proto-R152 exhibited greater diversity and less redundancy than the ones used by proto-D161 and proto-V19.

Second, to evaluate artifact localization, we examined whether the areas that the proto-R152 algorithm compares (the focus of “attention”) between an input slice and associated top-prototypes (prototypes with the highest similarity scores to the input slices) appeared relevant. We selected 100 2D slices at random from the original training set of 62 Class 1 scans from ABIDE 1, and examined the top 5 prototypes and the associated attention maps. One rater - Melanie Garcia - estimated that 52.4% of the attention maps were visually meaningful, in that artifacts were visible on the 2D image. For the remaining maps, either the artifact appeared elsewhere in the slice, or no obvious artifact could be detected by eye. Two examples of such attention maps are provided in **Figures S10 and S11.** This outcome suggests that while there is some congruence between human-identified and automatically identified artifacts, the algorithm may detect and rely on information that is not visible to the human eye. Future work will evaluate the attention maps and performance at the local scale in greater detail.

### 3.7 BIDS Docker app

We developed a BIDS-app (Gorgolewski et al., 2017) to share our model with the neuroimaging community. It is available on the open-source platforms GitHub and DockerHub. The model and instructions are available at: https://github.com/garciaml/BrainQCNet. The GPU/CUDA version is optimal. The average time to process a 3D sMRI scan using was about 1 minute 30 seconds on a laptop with one GPU Nvidia GEFORCE GTX 1060 (6GB memory) and 50 seconds on a machine with one GPU Nvidia RTX 3090 (24GB memory). While we strongly recommend the GPU version, there is also a CPU version available. Runtime will depend on the architecture available; in our experience, the average time to process a scan was about 30 minutes on a laptop with Intel Core I7-7700HQ processor (16GB memory), while it took about 10 minutes on an Intel Core i9-10850K (64GB memory).

## 4. Discussion

In this age of “big data’’, manual quality control of T1-weighted MRI scans is a time-consuming task requiring substantial experience and training. Our goal was to further advance the automatic detection of artifacts in sMRI scans by increasing the efficiency of the process. We trained an attention Deep Learning algorithm, ProtoPNet, paired with several different CNN architectures for the convolutional layer, to classify *minimally preprocessed* sMRI scans as pass/good quality and fail/poor quality. Specifically, the algorithms yielded class (0/1) predictions at the level of 2D image slices. These were converted to a probability value for each scan by computing the proportion of slices classified as fail/poor quality. Binary pass/fail global scan-level predictions were then generated by applying a threshold of 50% to the probability values. We evaluated our models’ performance by comparison to a reference tool in neuroscience (MRIQC) and to three traditional (non-attention) CNN models. Training, validation, and test sets comprised 4598, largely openly available sMRI scans from a large number of data collection sites, enabling the validation of the best-performed model using fully independent data.

Across convolutional layer architectures, the attention model ProtoPNet combined with a ResNet152 CNN architecture and trained on 10 epochs showed the best performance. On the first, non-independent, testing set (908 scans from ABIDE 1; Di Martino et al., 2014), this model performed equally as well as the reference tool, MRIQC (accuracy for high quality scans: 95.27% vs 91.1% for MRIQC; accuracy for medium and low quality scans: 47.89% vs 41.58% for MRIQC). Proto-R152 was also more sensitive than traditional CNNs, although less specific. On the second, independent, testing set (2141 scans from ABCD; Volkow et al., 2018; Karcher and Barch, 2021), the model showed excellent (91.4%) accuracy for low quality scans (i.e. high sensitivity). For high-quality scans, our model showed good prediction accuracy (82.4%), but this was lower than that of comparison models, including MRIQC (90.4%) and the CNN baseline models (from 99.2% to 99.9%). When we examined this more closely, we found that scans with a prediction falling in the mid-range of probabilities [0.5; 0.6] contained a mixture of good quality scans and moderately corrupted scans with more localized artifacts. If this “borderline” range was excluded, our model exhibited excellent accuracy for both pass and fail classes (accuracy for pass scans: 96.4%; accuracy for fail scans: 92.2%).

These data illustrate an advantage of our model - the ability to adjust global classification thresholds, or to isolate scans with probabilities falling within a specific range for further quality assessment. These parameters can be adjusted to make the classification categories more or less inclusive according to study needs. For applications where large samples are available and very high quality (artifact-free) data are required (e.g., computation of cortical thickness), the conservative 0.5 threshold could be retained. In other words, all the scans with a returned probability higher than 0.5 could be ruled out. This would have the disadvantage of removing some relatively good quality scans but the advantage of ruling out a greater proportion of lower quality scans than any other automatic method. If, on the other hand, a researcher had a smaller sample and less stringent quality requirements, a more liberal threshold of 0.6 could be set. This would mean that some scans with low severity or localized artifacts would be included in the study, but would offer the advantage that no good quality scans would be unduly eliminated. A third possibility is for researchers to retain all scans that have a global probability lower than 0.5, and to run one of our CNN models (or to manually evaluate or run MRIQC) on scans that have a global probability between 0.5 and 0.6 to separate the good from moderately corrupted scans. To facilitate these possibilities, our BIDS-app (https://github.com/garciaml/BrainQCNet) outputs a CSV file containing probability scores for each scan.

Our study demonstrates that Deep Learning is a promising method for increasing the speed of scan quality evaluation by reducing the computational time required, without compromising classification accuracy. Importantly, preprocessing was minimal - avoiding even the need for data reorienting, since our model was trained to process transformed (rotated, skewed, sheared) 2D image slices from the three axes (sagittal, coronal, axial). This simplifies the process compared to approaches where knowing the data orientation is necessary (Sujit et al., 2019). To generate a global prediction for a single 3D scan on a GPU machine, our model currently takes 1 minute to process one scan (50 seconds on a machine with one GPU Nvidia RTX 3090, 24GB memory; 1 minute 30 seconds on a laptop with one GPU Nvidia GEFORCE GTX 1060, 6GB memory). On a CPU machine, our model is slower but still relatively fast (10 minutes on an Intel Core i9-10850K; 64GB memory; 30 minutes on an Intel Core I7-7700HQ processor, 16GB memory). We have openly shared our code so it can be further adapted to other architectures.

In order to save resources and encourage sustainable practices, we have also shared the global scores predicted by our best model for the scans we used from ABIDE 1 and 2 (Di Martino et al., 2014; Di Martino et al., 2017), ADHD200 (Bellec et al., 2017) and ABCD (Volkow et al., 2018; Karcher and Barch, 2021). The scores are available through our GitHub repository: https://github.com/garciaml/BrainQCNet_paper_results. In addition, we have shared a version of the app containing the traditional (non-attention) CNN models. Even though our data showed that these algorithms are less sensitive (have a greater number of false negatives), they nonetheless show excellent accuracy (true negatives) for good quality (pass) scans. These characteristics may be of use for certain applications or may offer possibilities for further refinement.

Deep Learning models often lack interpretability - attention models reflect an attempt to address this. As implemented here, the attention ProtoPNet model enables the localisation of regions in the input images that contribute significantly to classification. This might help to identify specific brain regions that are more vulnerable to artifacts, such as motion, or highlight a scanner quality issue that can be addressed to avoid future data loss. We have made it easy to inspect regions exhibiting local artifacts using our BIDS-app, using the parameter “n_area.” Details on how to do this can be found in the documentation.

Future work will focus on improving our algorithm by running further experiments with other CNN-bases, such as ResNet34 or DenseNet121, and examining the effects of prototype selection. In addition, we plan to increase the training set, as well as the variety of artifacts in the set of prototypes, since our approach was not exhaustive. It is likely that signals in the background are leveraged by the current attention algorithm and this behavior should be studied more precisely. To take a wider view, it is clear that MRI scan quality is a continuous spectrum; pass/fail (good/bad) thresholds can seem arbitrary and simplistically binary. Scan quality would be better captured by a more sophisticated label, but this is very difficult to implement concretely. Future work should investigate whether such a prediction can be obtained by incorporating additional information about the location/extent of artifact.

Investigating whether our approach could be applied to other MRI modalities is another important future direction. Quality Control of functional MRI is a considerable challenge that is exacerbated by the advent of Big Data. Future work will examine whether our approach can be adapted for data with a temporal dimension so that it could be applied to fMRI data in a framewise manner to enable faster and automated data quality control.

There is further scope for improvement of our algorithm and app - particularly in terms of processing speed. While the model already exhibits fast performance on GPU, we have not yet attempted to optimize the implementation by better distributing the computations or better use of infrastructure types. These possibilities will be investigated for future versions of the app, to further foster reusability.

Finally, to our knowledge, our BIDS-app is the first app that applies Deep Learning to neuroimaging and is built to be used on CUDA GPU machines. By sharing our code, we are providing the community with a new BIDS-app template for Deep Learning applications, facilitating the sharing of Deep Learning models in the community and helping to maximize reproducibility and collaboration.

## 5. Conclusions

In this work, we introduced a novel Deep Learning approach for the automatic evaluation of the quality of minimally preprocessed structural MRI scans. Our method is scalable to big datasets by taking advantage of new technologies like GPU machines with high-computing capacity. Paths to improve our model include incorporating additional CNN architectures and manually selecting the prototypes used by the model to increase the diversity of artifacts represented during training. Our approach could be further adapted to functional MRI, as well as to other types of MRI scans and organs. Our model is already freely available for use and development by the community via the app BrainQCNet (https://github.com/garciaml/BrainQCNet). Since all our code is open-source, the app can be used as a template for future applications of Deep Learning in neuroimaging.

## 6. Acknowledgements and Funding

This research was supported by the Hypercube Institute (Paris, France), a Trinity College Dublin Postgraduate Research Studentship (1252) awarded by the School of Medicine, and an Irish Research Council Government of Ireland Postgraduate Scholarship award (GOIPG/2021/1508).

## 7. Disclosure of competing interests

None.

### Terms and abbreviations

- **CNN**: Convolutional Neural Networks, a category of Deep Learning algorithm
- **ML**: Machine Learning
- **DL**: Deep Learning
- **Epoch**: a hyperparameter that defines the number of times that the learning algorithm has optimized the parameters on the entire training dataset.
- **ProtoPNet**: Prototypical Part Network model
- **VGG19**: Visual Geometry Group model, a type of very deep convolutional neural network with 19 layers in the model;
- **ResNet152**: Residual Networks model with 152 layers
- **DenseNet161**: Densely Connected Convolutional Networks with 161 layers
- **proto-V19:** ProtoPNet model with a VGG19 architecture in the CNN part
- **proto-R152:** ProtoPNet model with a ResNet152 architecture in the CNN part
- **proto-D161:** ProtoPNet model with a DenseNet161 architecture in the CNN part

## Notes

### Competing Interest Statement

The authors have declared no competing interest.

### Summary of Updates

To address issues, we have: - Thoroughly revised the manuscript text, paying particular attention to clarity of expression and argument. Revised text appears in blue. - Expanded the supporting content (Docker apps, code, and documentation) available through our GitHub repo (https://github.com/garciaml/BrainQCNet), so that our tool is more accessible and usable on both GPU and CPU machines. We have created and shared two demo videos (via YouTube) to help people who would like to run our BIDS-app on CPU machines and on NVIDIA GPU machines with CUDA installed. We have also shared non-containerized code for CPU machines that can be readapted and optimized locally. Finally, we also shared all the CNN models we trained as baseline comparisons, and we built a BIDS-app that uses them. - Revised our analyses and results to include an examination of specificity (rate of True Negatives rate) and sensitivity (rate of True Positives). In addition, we trained three CNN models to provide additional baseline comparisons for our study. We showed that our attention model had superior sensitivity (True Positives, i.e., successful identification of corrupted/poor quality scans), which is an important advantage. - Revised our discussion of computational efficiency and resource use, tempering any comparisons with other tools. - Expanded reference to previous projects that used Deep Learning models to detect the quality of sMRI scans. We have contextualized our work with reference to these approaches and explained why we chose to train an attention model over a traditional CNN. - Performed additional analyses and discussed in greater depth the interpretation of our model at a local scale. For future versions of our tool, we will focus on strategies to improve the local detection of artifacts.

