## Supplemental Information for "BrainQCNet: a Deep Learning attention-based model for the automated detection of artifacts in brain structural MRI scans"

### 9. Supplemental Information

#### 9.1 Comparison of the distribution of probabilities between models

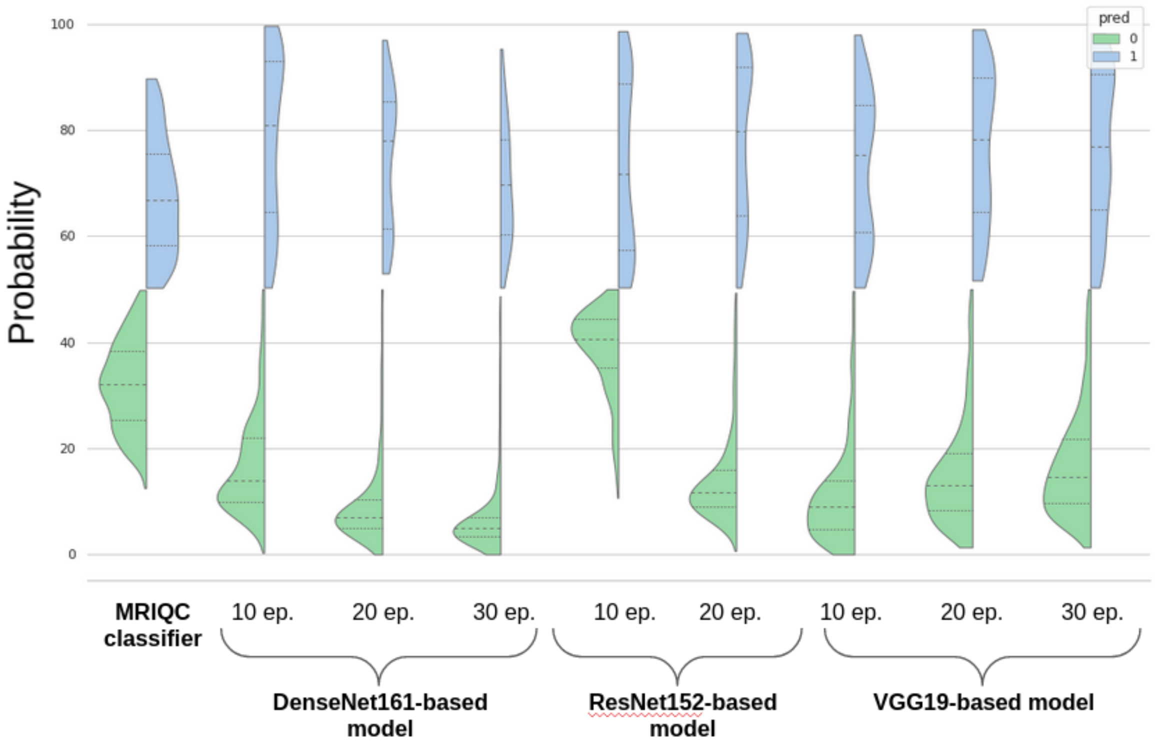


**Figure S1.** Comparison of the distribution of probabilities for the test set (908 scans), colored by predicted class: green for Class 0 (good quality scans), blue for Class 1 (medium/low quality scans).

In **Figure S1**, we can see that the distribution of predictions of Class 0 scans (green) looks gaussian for our models. In contrast, the distribution of predictions for Class 1 (blue) looks like a gaussian mixture. This distribution shape is expected since there are some scans that are globally corrupted scans, but others with only local artifact, or less severe global artifact. The proportion of slices classified as Class 1 will therefore be different for the two types.

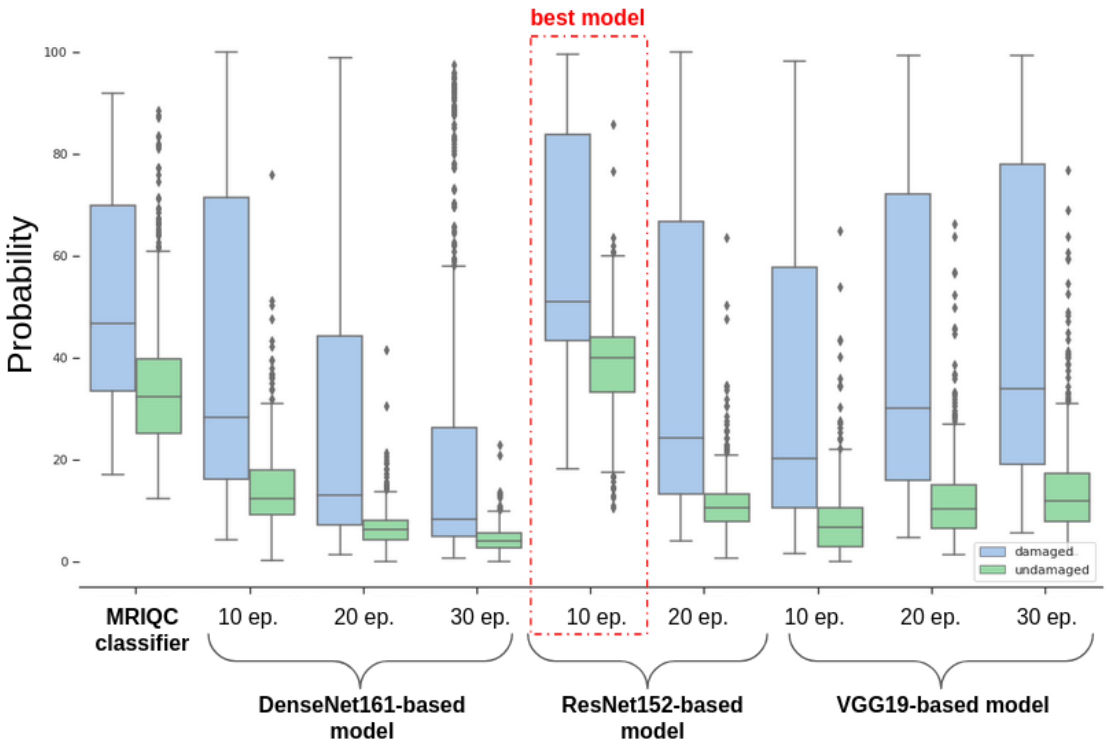


**Figure S2.** The boxplots show the predicted probabilities (% slices predicted to be Class 1, poor quality) for scans manually judged to be free from artifact (Class 0 - good quality; green) vs. those manually judged to be contain some artifact (Class 1 - blue) for all models and for MRIQC, using 980 scans from ABIDE 1. The figure shows that there is some overlap in the global probabilities for Class 0 and Class 1 scans, although this varies by model and by epoch. The greater the overlap, the more False Positives and False Negatives there are. The overlap is least, and optimally located (around 40% probability) for the best-performing model, **proto-R152 (10 epochs).**

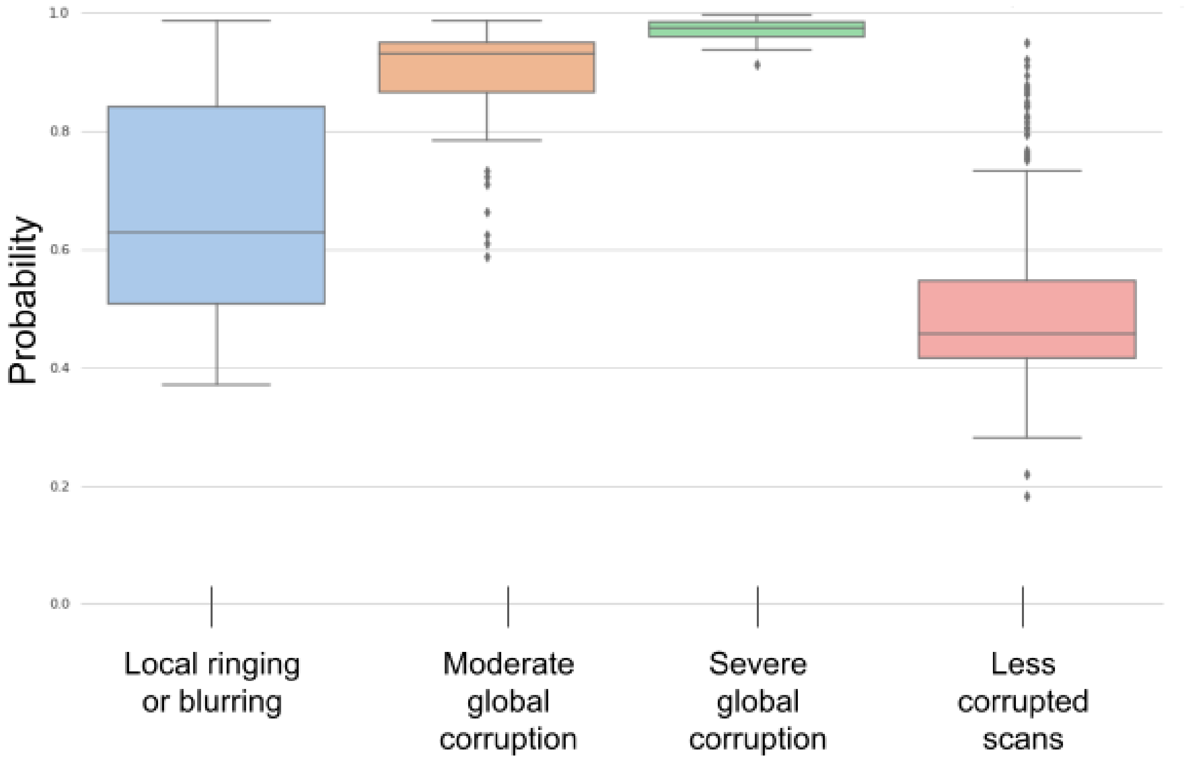


**Figure S3.** Comparison of probabilities for global predictions from proto-ResNet152 trained on 10 epochs for 416 Class 1 (poor quality) scans from ABIDE 1 (30 very low quality scans included in the training set, 6 very low quality scans included in the validation set, 380 less severely poor quality scans included in the test set). 51 scans have local ringing or blurring (blue), 60 are globally corrupted but medium quality (orange), 36 are globally corrupted and very low quality (green; i.e. score=4,4,4,4 and artifacts present on all the 2D slices), 269 are less severely corrupted or exhibit localized artifact only (red).

**Figure S3** shows that each of the categories of artifact severity are well segregated in terms of their predicted probabilities: globally corrupted scans have probabilities very close to 1, while scans with moderate artifact (e.g., ringing or blurring) have probabilities spread between 0.5-0.8, and other scans with localized or less severe artifact have probabilities around 0.5.

The classes (Local/Severe/Moderate/Less corruption) were defined as follows:

- Severe/Moderate/Less: refers to an artifact or a set of artifacts that was globally evident on the scan. Severe means that the scan was highly corrupted: at least one of the four artifact scores (blurring; ringing; CNR WM/GM; CNR subcortical structures) was 4. Moderate means that at least one of the four artifact scores was 3. Less means that at least one of the four artifact scores was 2.
- Local: refers to an artifact or a set of artifacts (with scores between 2 and 4) that was present only on a demarcated area of the scan, and on for less than half of the slices.

We also evaluated the results on slices for the 66 scans from ABIDE 1 we annotated with local ringing and/or local blurring. We found that in the extremities, the algorithm tends to predict the slices as Class 1, even in the cases it should be Class 0. This means that slices near the edge of the field-of-view containing few brain tend to be identified as corrupted by the algorithm. This might explain why the global distribution of probabilities of the model proto-ResNet152-10ep is higher than the ones of other models (see **Figures S1 and S2**).

We also found an axis effect at the accuracy level (no effect between the distribution of scores) - while predictions for sagittal images were 89.3% accurate, accuracy for coronal images were 86.4% accurate, and for axial views, 78.8% accurate.

#### 9.2 Multi-site effect

| | good quality - 528 scans | globally medium corrupted - 60 scans | local ringing or blurring - 51 scans | other less corrupted scans - 269 scans |
|---|---|---|---|---|
| **CALTECH** | accuracy: 1.0<br>n scans: 34 | na | na | accuracy: 0.0<br>n scans: 2 |
| **CMU** | accuracy: 1.0<br>n scans: 24 | na | na | accuracy: 0.3333<br>n scans: 3 |
| **KKI** | accuracy: 1.0<br>n scans: 25 | accuracy: 1.0<br>*n scans:* 3 | na | accuracy: 0.5714<br>n scans: 14 |
| **LEUVEN_1** | accuracy: 0.9259<br>n scans: 27 | na | na | accuracy: 0.5<br>n scans: 2 |
| **LEUVEN_2** | accuracy: 0.9565<br>n scans: 23 | na | accuracy: 1.0<br>n scans: 1 | accuracy: 0.2<br>n scans: 10 |
| **MAX_MUN** | accuracy: 0.9286<br>n scans: 28 | accuracy: 1.0<br>n scans: 2 | accuracy: 1.0<br>n scans: 1 | accuracy: 0.8<br>n scans: 10 |
| **NYU** | accuracy: 0.9146<br>n scans: 82 | accuracy: 1.0<br>n scans: 1 | accuracy: 0.5882<br>n scans: 17 | accuracy: 0.2714<br>n scans: 70 |
| **OHSU** | accuracy: 0.9091<br>n scans: 22 | accuracy: 1.0<br>n scans: 1 | na | na |
| **OLIN** | accuracy: 0.75<br>n scans: 12 | na | accuracy: 1.0<br>n scans: 2 | accuracy: 0.4286<br>n scans: 7 |
| **PITT** | accuracy: 0.9524<br>n scans: 21 | na | accuracy: 1.0<br>n scans: 5 | accuracy: 0.3913<br>n scans: 23 |
| **SBL** | accuracy: 1.0<br>n scans: 26 | na | na | accuracy: 0.0<br>n scans: 4 |
| **SDSU** | accuracy: 0.8<br>n scans: 10 | accuracy: 1.0<br>n scans: 10 | na | accuracy: 0.8<br>n scans: 10 |
| **STANFORD** | na | accuracy: 1.0<br>n scans: 10 | accuracy: 0.8333<br>n scans: 12 | accuracy: 0.6667<br>n scans: 6 |
| **TRINITY** | accuracy: 1.0<br>n scans: 34 | accuracy: 1.0<br>n scans: 3 | accuracy: 1.0<br>n scans: 1 | accuracy: 0.0<br>n scans: 7 |
| **UCLA_1** | accuracy: 0.8958<br>n scans: 48 | accuracy: 1.0<br>n scans: 6 | accuracy: 0.6667<br>n scans: 3 | accuracy: 0.8<br>n scans: 5 |
| **UCLA_2** | accuracy: 1.0<br>n scans: 7 | accuracy: 1.0<br>n scans: 3 | accuracy: 1.0<br>n scans: 1 | accuracy: 0.4286<br>n scans: 7 |
| **UM_1** | accuracy: 1.0<br>n scans: 27 | accuracy: 1.0<br>n scans: 7 | accuracy: 0.8<br>n scans: 10 | accuracy: 0.1471<br>n scans: 34 |
| **UM_2** | accuracy: 1.0<br>n scans: 13 | na | accuracy: 0.6667<br>n scans: 3 | accuracy: 0.25<br>n scans: 12 |
| **USM** | accuracy: 1.0 | na | na | accuracy: 0.25 |

| | | | | |
|---|---|---|---|---|
| | n scans: 60 | | | n scans: 4 |
| **YALE** | accuracy: 1.0<br>n scans: 5 | accuracy: 1.0<br>n scans: 5 | accuracy: 0.5<br>n scans: 4 | accuracy: 0.1795<br>n scans: 39 |

**Table S1.** Predictions for each data collection site in the test set (908 scans) for proto-ResNet152 trained on 10 epochs.

**Table S1** displays the predictions of the model proto-ResNet152 trained on 10 epochs for each data collection site in the first test set of 908 scans from the ABIDE 1 dataset.

For Class 0 (good quality, pass) scans, the model attained 100% accuracy for 10 out of 19 sites, >90% for 16 of 19, and accuracy of at least 75% for the remaining 3 sites. Overall the mean accuracy across sites is 94.9% with a standard deviation of 0.07. There does not appear to be a significant effect on site effect on the prediction of the good quality (Class 0) scans.

For Class 1 (poor quality, fail) scans with moderate levels of artifact, accuracy is 100% across sites. For scans with less severe or localized artifact, there is more variability across sites, but there is also large variation in the number of scans in that category, so it is difficult to quantitatively assess whether there is a significant site effect.

#### 9.3 Prototypes mostly used by proto-R152

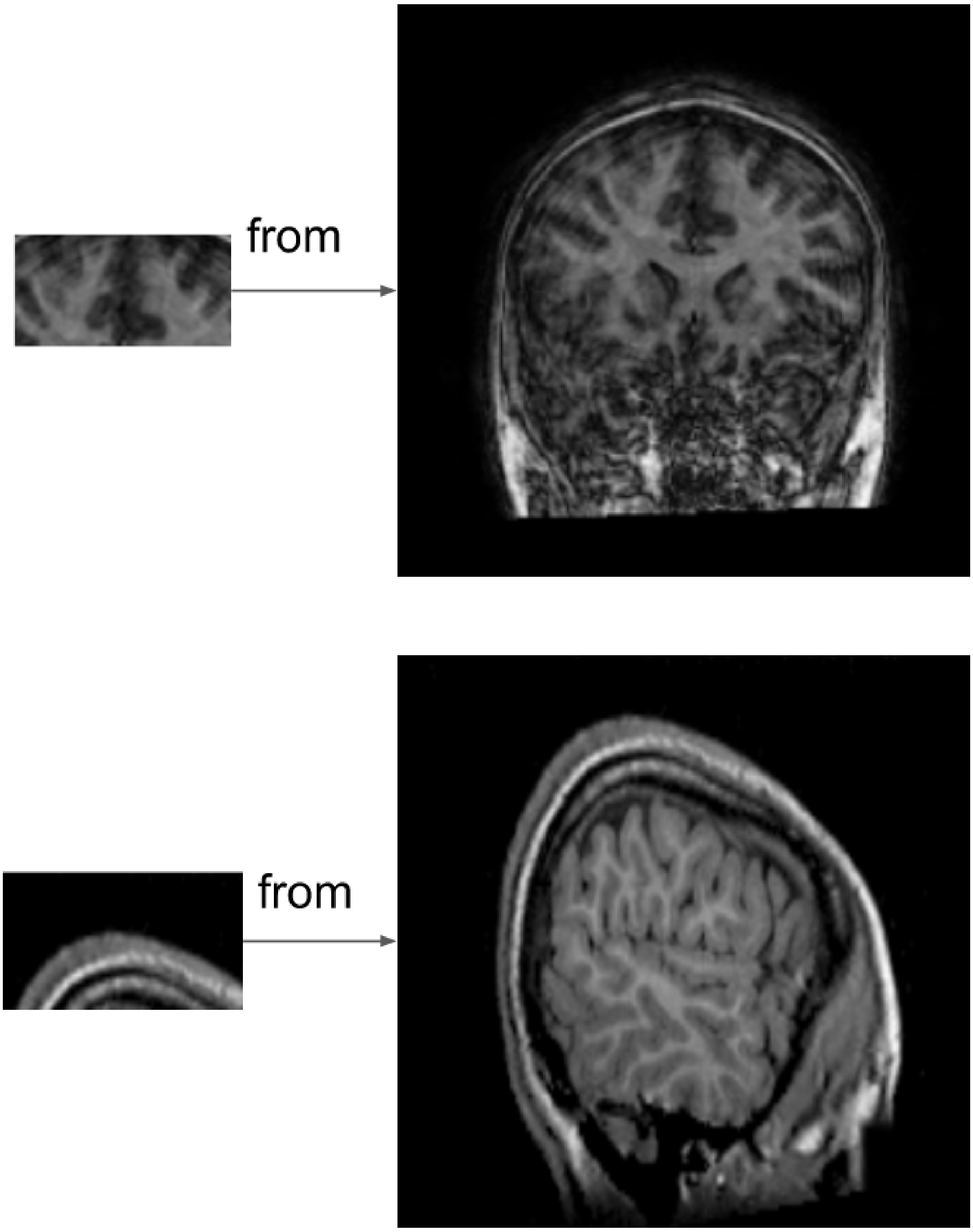


**Figure S4.** Prototypes that commonly appeared in the top-5 prototypes (i.e., those prototypes with the top 5 highest ranking similarity scores to the input 2D slices).

#### 9.4 Examples of activation maps

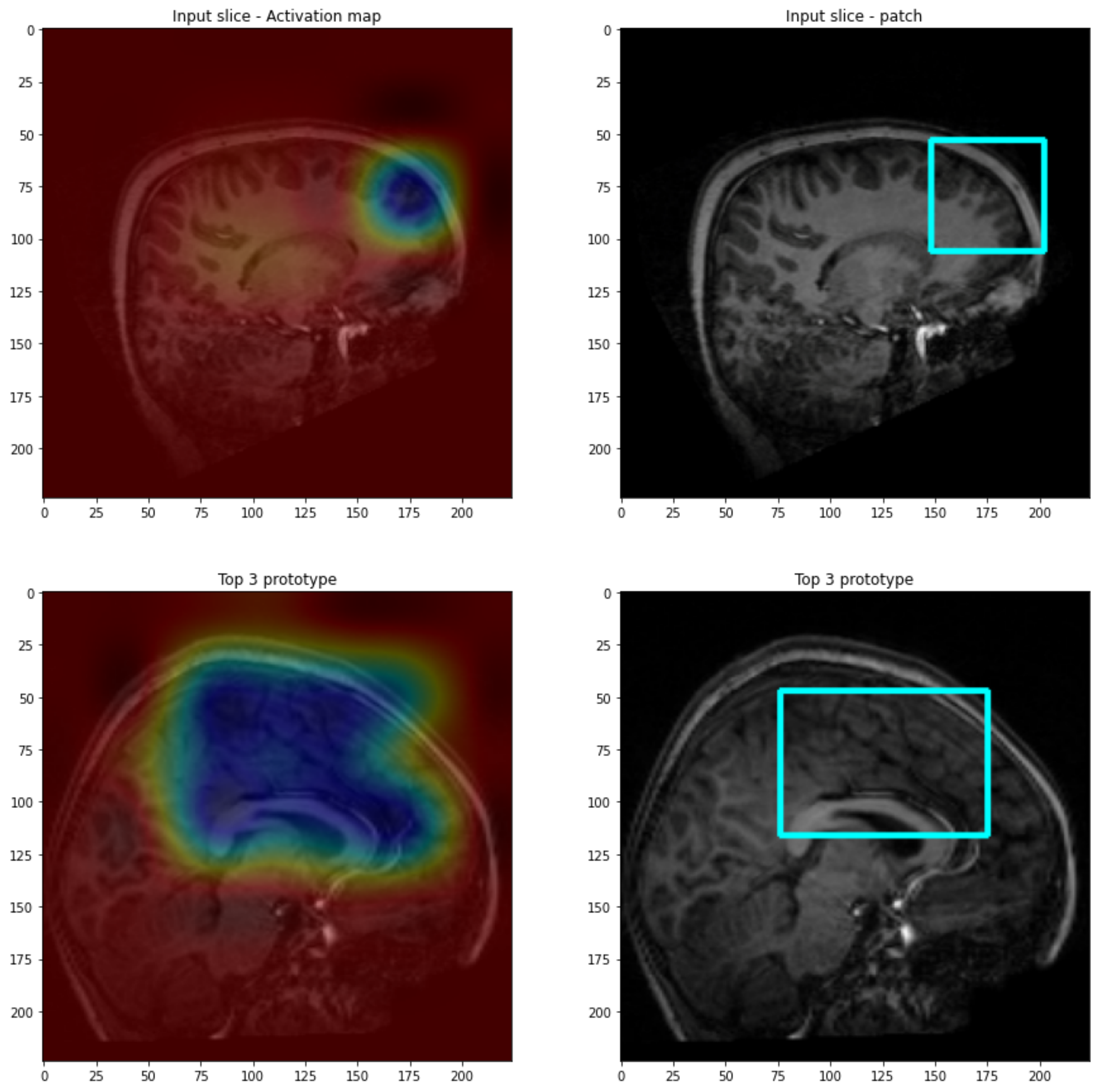


**Figure S5.** Examples of artifact maps and prototypes judged to be meaningful: the upper panel shows the input slice, the lower panel shows the top-3 prototype for the model proto-R152 trained on 10 epochs (i.e., the prototype with the third highest ranking similarity score to the input 2D slices).

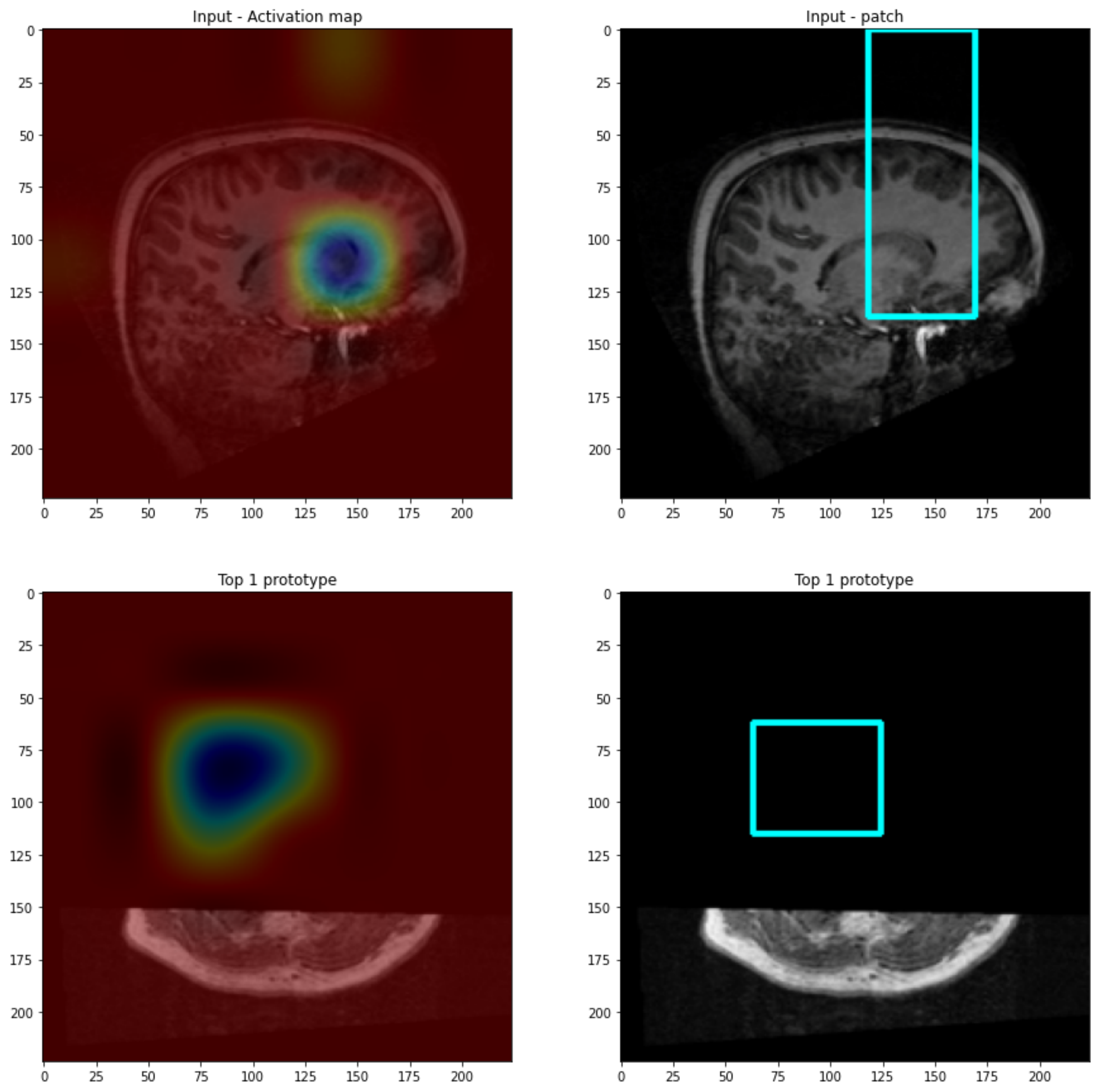


**Figure S6.** Examples of artifact maps and prototypes judged not to be meaningful: the upper panel shows the input slice, the lower panel shows the top-1 prototype for the model proto-V19 trained on 30 epochs (i.e., the prototypes with the highest ranking similarity score to the input 2D slices).

##### 9.5 Mann-Whitney U-tests between the predicted scores of QC categories of the ABCD data

| variable 1 | variable 2 | U-val | alternative | p-val | RBC | CLES |
|---|---|---|---|---|---|---|
| **pass** | **fail** | 13814,5 | two-sided | $1{,}844.10e^{-93}$ | 0,899 | 0,051 |
| **pass** | **questionable** | 208800,0 | two-sided | $1{,}183.10e^{-56}$ | 0,459 | 0,271 |
| **fail** | **questionable** | 94121,5 | two-sided | $1{,}004.10e^{-48}$ | -0,701 | 0,850 |

**Table S2.** Mann-Whitney U-tests between the scores returned by BrainQCNet on each QC category of the ABCD dataset: pass, questionable, fail. It shows that the distribution of scores are well distinct between each other, which is an expected behavior for our QC algorithm.
